# INPP5D/SHIP1 regulates inflammasome activation in human microglia

**DOI:** 10.1101/2023.02.25.530025

**Authors:** Vicky Chou, Seeley B. Fancher, Richard V. Pearse, Hyo Lee, Matti Lam, Nicholas T. Seyfried, David A. Bennett, Phillip L. De Jager, Vilas Menon, Tracy L. Young-Pearse

**Author notes:** Correspondence to: Ann Romney Center for Neurologic Diseases, Brigham and Women’s Hospital and Harvard Medical School, 60 Fenwood Rd, Boston, MA, 02115, E-mail address (T. Young-Pearse).

## Abstract

Microglia and neuroinflammation are implicated in the development and progression of Alzheimer’s disease (AD). To better understand microglia-mediated processes in AD, we studied the function of INPP5D/SHIP1, a gene linked to AD through GWAS. Immunostaining and single nucleus RNA sequencing confirmed that INPP5D expression in the adult human brain is largely restricted to microglia. Examination of prefrontal cortex across a large cohort revealed reduced full length INPP5D protein levels in AD patient brains compared to cognitively normal controls. The functional consequences of reduced INPP5D activity were evaluated in human induced pluripotent stem cell derived microglia (iMGLs), using both pharmacological inhibition of the phosphatase activity of INPP5D and genetic reduction in copy number. Unbiased transcriptional and proteomic profiling of iMGLs suggested an upregulation of innate immune signaling pathways, lower levels of scavenger receptors, and altered inflammasome signaling with INPP5D reduction. INPP5D inhibition induced the secretion of IL-1ß and IL-18, further implicating inflammasome activation. Inflammasome activation was confirmed through visualization of inflammasome formation through ASC immunostaining in INPP5D-inhibited iMGLs, increased cleaved caspase-1 and through rescue of elevated IL-1ß and IL-18 with caspase-1 and NLRP3 inhibitors. This work implicates INPP5D as a regulator of inflammasome signaling in human microglia.

## INTRODUCTION

There is an increasing focus on understanding the role of microglia in neurodegenerative disorders including late-onset Alzheimer’s disease (LOAD), a disease defined by the accumulation of amyloid beta rich plaques and neurofibrillary tangles containing tau. Microglia are proposed to play a variety of critical roles during disease progression including synaptic engulfment^1^, cytokine release, and phagocytosis of amyloid beta^2^. Further, recent GWAS studies have implicated innate immune processes in LOAD, supporting the importance of understanding the neuroimmunological processes underlying the risk and progression of AD.

The inflammasome is a multimeric complex involved in innate immune signaling that is induced in response to inflammatory insult^3^. Inflammasome activation involves conformational changes in NLR family members that allow for association and oligomerization with the adaptor protein ASC (PYCARD), and inactive pro-caspase-1 (CASP1). Upon inflammasome formation, pro-CASP1 is cleaved to generate active CASP1 which in turn cleaves pro-IL-1ß and pro-IL-18 resulting in extracellular release of these cytokines^3^. Inflammasome activity is associated with several inflammatory disorders and more recently with AD^4–6^. In murine models of familial AD, inflammasome activation in microglia has been shown to contribute to amyloid beta and tau pathology^4,5^. The regulation of inflammasome activation in human microglia is poorly understood, thus the identification of other members of this signaling cascade may prove valuable towards targeting inflammasome activity in disease.

SNPs at the *INPP5D* locus have been associated with LOAD through GWAS^7–10^. The lead SNPs, rs35349669, rs10933431, and rs7597763, are each located within introns of INPP5D, and it is unclear if these SNPs are associated with altered RNA or protein levels of INPP5D in brain tissue. INPP5D was first characterized through loss-of-function mutations that cause leukemia and other related myeloproliferative cancers^11^. Subsequent studies revealed that lack of INPP5D expression in peripheral macrophages resulted in dysregulation of immune reactivity^12^. INPP5D encodes a 145 kDA membrane-associated phosphatase that acts as a negative regulator of PI3K/Akt signaling by hydrolyzing the 5’-phosphate of the secondary messenger, phosphatidylinositol (3,4,5)P_3_ to generate phosphatidylinositol (3,4)P_213_. Elevated INPP5D activity results in reduced levels of phosphorylated Akt, which in turn effects cell metabolism and survival signaling^14^. INPP5D knockout mouse models experience a peripheral over-proliferation of myeloid cells that results in early postnatal lethality^12^. In addition, INPP5D has been reported to function as a scaffolding protein, binding with DNAX-activating protein of 12 kDa (DAP12) to prevent PI3K association with triggering receptor expressed on myeloid cells 2 (TREM2)^15^. INPP5D also influences phagocytic activity of macrophages through binding to FcgammaR and ROS production through interactions with Dectin-1/CLEC7A^16,17^.

Understanding the role of INPP5D in microglia offers insight into the dynamic regulation of microglia in the non-disease state and potentially advances our understanding of AD pathogenesis. Here, we show that INPP5D expression is largely restricted to microglial cells in the human adult brain and confirm previous findings that RNA levels of INPP5D are elevated in AD brain^18^ However, through quantitative western blotting, we show that protein levels of full length, water-soluble INPP5D are *reduced* in AD brain. To study the functional consequences of reduced INPP5D, we use both pharmacological inhibition and CRISPR-Cas9 genome engineering to reduce INPP5D activity in human iPSC-derived microglia. Through unbiased RNA and proteomic profiling and a series of pharmacological manipulations, we demonstrate that reduction of INPP5D activity induces changes in immune signaling and, more specifically, the activation of the inflammasome.

## RESULTS

### INPP5D expression is restricted to human microglia and reduced in Alzheimer’s disease

We first confirmed INPP5D expression in human microglia using immunostaining. Human post-mortem brain tissue was fixed, cryosectioned, and immunostained for INPP5D and IBA1, a microglial and macrophage-specific calcium-binding protein. INPP5D and IBA1 co-localized to the same subset of cells, suggesting that INPP5D is expressed in microglia in the human brain (**Fig. 1a**).

**Figure 1:**
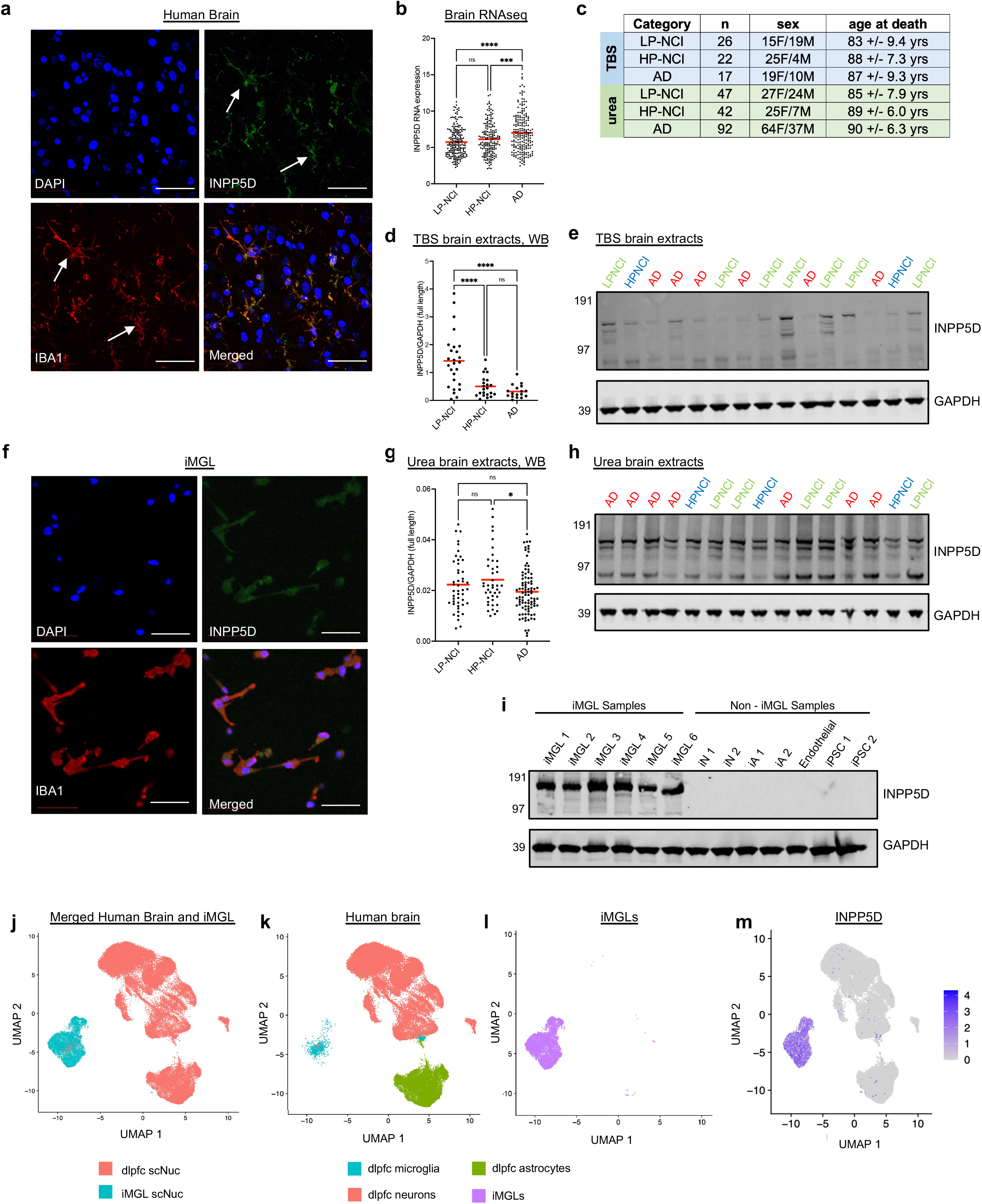
INPP5D expression in the human brain is restricted to microglia and reduced in Alzheimer’s disease. **a**. Human brain sections (25 microns) immunostained with IBA1 and INPP5D. DNA is stained with DAPI. Imaged using confocal microscopy. Scale bars = 50 um. **b**. Relative levels of INPP5D RNA as determined by RNA sequencing of 584 ROSMAP human dorsolateral prefrontal cortex samples (198 LP-NCI, 178 HP-NCI, 208 AD; source data from^22^). One-way ANOVA with Tukey’s multiple comparisons test. **c**. A summary table detailing the brain samples assayed in Fig. 1d-e, 1g-h. **d**. Western blot quantification of INPP5D and GAPDH in TBS postmortem brain (mPFC) brain extracts from LP-NCI, HP-NCI, and AD patients. One-way ANOVA with Tukey’s multiple comparisons test. **e**. Representative western blot of TBS extracts from postmortem human brain for INPP5D and GAPDH. **f**. iMGLs immunostained with IBA and INPP5D. DNA is stained with DAPI. Imaged using confocal microscopy. Scale bars = 50 um. **g**. Western blot quantification of INPP5D and GAPDH from urea extracts from 181 ROSMAP human brain samples. One-way ANOVA with Tukey’s multiple comparisons test. Outliers were eliminated with ROUT Q=10%. **h**. Representative western blot of urea extracts from postmortem human brain for INPP5D and GAPDH levels^23^. **i**. Western blot of INPP5D and GAPDH in iMGLs, iPSC-derived neurons (iN), iPSC-derived astrocytes (iA), iPSC-derived endothelial cells, and undifferentiated iPSCs. **j**. UMAP plots of sNucRNAseq of iMGLs combined with sNucRNAseq of dorsolateral prefrontal cortex (dlpfc) from 12 human postmortem brain samples^31^ using Harmony to integrate across data sets. Single nucleus samples from human brain and iMGLs are depicted separately in **k** and **l**. 7121 iMGL nuclei, 55,671 nuclei from the postmortem human brain (39,239 glutamatergic neurons, 14,958 astrocytes, 1,474 microglia) **m**. Relative INPP5D expression across the iMGL and human brain single nucleus samples shown in **j**. For b,d,g: ns = not significant, * p < 0.05, ***p < 0.001, **** p < 0.0001

Next, we examined human brain tissue from the Religious Order Study (ROS) and Rush Memory and Aging Project (MAP) for RNA and protein expression of INPP5D^19–23^. We compared INPP5D levels across three categories, low pathology, not cognitively impaired (LP-NCI), high pathology, not cognitively impaired (HP-NCI) and AD. LP-NCI individuals did not have a clinical or pathological diagnosis of AD. HP-NCI had a pathological diagnosis of AD but no cognitive impairment, and “AD” had both clinical and pathological diagnoses of AD. Brain samples were age and sex matched. At the RNA level, we queried a published dataset^24^ to confirm previous reports^18^ that cohorts of AD brain tissue expresses higher levels of INPP5D compared to NCI brain tissue (**Fig. 1b**). INPP5D is a cytosolic protein with a predicted molecular weight in its full-length form of 145 kDa. However, multiple splice variants have been reported, some of which encode forms of INPP5D that do not contain the phosphatase domain and are predicted to be inactive^25^. To quantify protein levels of full length INPP5D (>100 kDa), we obtained medial prefrontal cortex (mPFC) samples from a cohort of 92 individuals (**Fig 1c, Supplemental Table 1**). We homogenized these in tris buffered saline (TBS) to extract soluble, cytosolic proteins and performed Western blotting to quantify full length INPP5D. A significant reduction in INPP5D protein levels was observed in AD and HP-NCI groups compared to LP-NCI brain tissue (**Fig. 1d, 1e**). While INPP5D has been described to function in the cytosol, it may be partitioned into multiple subcellular compartments within microglia in the brain and TBS-soluble lysates may not capture the totality of full length INPP5D. Further, in addition to Aß and tau, a multitude of proteins accumulate in insoluble aggregates in the “high pathology” elderly brain. To quantify the entirety of full length INPP5D, we obtained a second cohort of brain tissue extracted with urea (**Supplemental Table 2, Fig 1c**). Quantification of full length INPP5D in urea-solubilized human brain tissue by western blot revealed lower levels of INPP5D in the population of AD brain tissues relative to LP-NCI individuals, with significantly lower levels in AD relative to HP-NCI (**Fig. 1g, 1h**). These data suggest that full length INPP5D is reduced in AD brain.

We next established a manipulable human experimental system for assaying INPP5D function using iPSC lines from ROSMAP individuals. This was accomplished by adapting an existing published protocol that generates iPSC-derived microglia-like cells (iMGLs) after 40 days of differentiation (**Extended Data 1a, 1b**)^26,27^. INPP5D expression in iMGLs was confirmed using immunocytochemistry with antibodies to INPP5D and IBA1 and the two signals co-localized to all (>99%) resultant cells (**Fig. 1f**). Using established protocols, we also generated iPSC-derived astrocytes (iAs)^28^, induced neurons (iNs)^29,30^, and iPSC-derived endothelial cells^31,32^ and evaluated INPP5D expression in each of these cell types using western blotting. INPP5D expression was restricted to iMGLs and INPP5D protein levels were consistent across iMGL differentiations (**Fig. 1i**).

To further confirm the utility of this model for studying INPP5D function, we performed single-nucleus RNA sequencing (sNucRNAseq) to compare the genetic signatures of the iMGLs to sNucRNAseq of human post-mortem frozen dorsolateral prefrontal cortex brain tissue from 12 adult individuals from the ROS-MAP studies (6 male and 6 female, ranging in age at death of 78-98 years, data from ^33^). Data from astrocytes, glutamatergic neurons, and microglia from brain tissue were merged and integrated with data from the iMGLs samples using Harmony^34^. The iMGLs cluster overlapped in UMAP space with the human microglia and not with neurons or astrocytes, suggesting transcriptional similarity between iMGLs and human brain microglia (**Fig. 1j-l**). Examination of *INPP5D* expression within these populations confirmed that expression is largely confined to the brain tissue microglia and iMGL cluster (**Fig. 1m, Extended Data Fig. 1c, d**). Within brain microglia, INPP5D is expressed throughout all subclusters of microglia, with no significant enrichment in any specific subtype of brain microglia (**Extended Data Fig. 1e**). Taken together, these data demonstrate that INPP5D expression is restricted to microglia in the human brain and INPP5D protein levels are reduced in AD patient brain tissue.

### Acute inhibition of INPP5D induces alterations of RNA and protein expression profiles associated with innate immune processes

As there are lower INPP5D protein levels in AD brain tissue when compared to controls, we interrogated a model of decreased INPP5D activity through transcriptomic and proteomic profiling following acute inhibition. Here, we used 3-alpha-Aminocholestane (3AC), a compound previously reported to be a selective inhibitor of INPP5D^35^, which has been utilized in several studies to reduce INPP5D activity in vivo in mice and in vitro cell culture ^35–43^. A recent study reported reduced INPP5D activity in primary mouse microglia treated with 3AC at a concentration (1.25 uM) well below its reported IC_50_ (10 uM)^35,41^. Here, iMGL cultures were treated with this low concentration for 6 hours. Cells were then collected for RNA sequencing and tandem mass tag mass spectrometry (MS). After filtering to remove weakly detected gene-level transcripts, 12,669 genes were quantified and of these 2,004 were differentially expressed between vehicle and 3AC treatment (**Fig. 2a**). 7,866 proteins were quantified by MS, and 159 differentially expressed proteins (DEPs) were identified comparing vehicle and 3AC treatment (**Fig. 2b**). Examination of overlap between RNA and protein changes revealed a set of genes that showed concordant changes in both RNA and protein datasets including CD33, a microglial gene associated with LOAD through GWAS, and FCER1G, a component of Fc receptors which has been reported to biochemically interact with INPP5D^44^ (**Fig. 2c**). Examination of AD-GWAS associated genes revealed 9 genes in addition to CD33 which were differentially expressed at the protein level: EIF4G3, TMEM106B, CTSH, SORL1, GRN, FDFT1, IL34, PTK2B, CD2AP (**Fig. 2d**). Of those genes, CD33 and PTK2B were both changed concordantly at the RNA and protein level (**Fig. 2e**).

**Figure 2.**
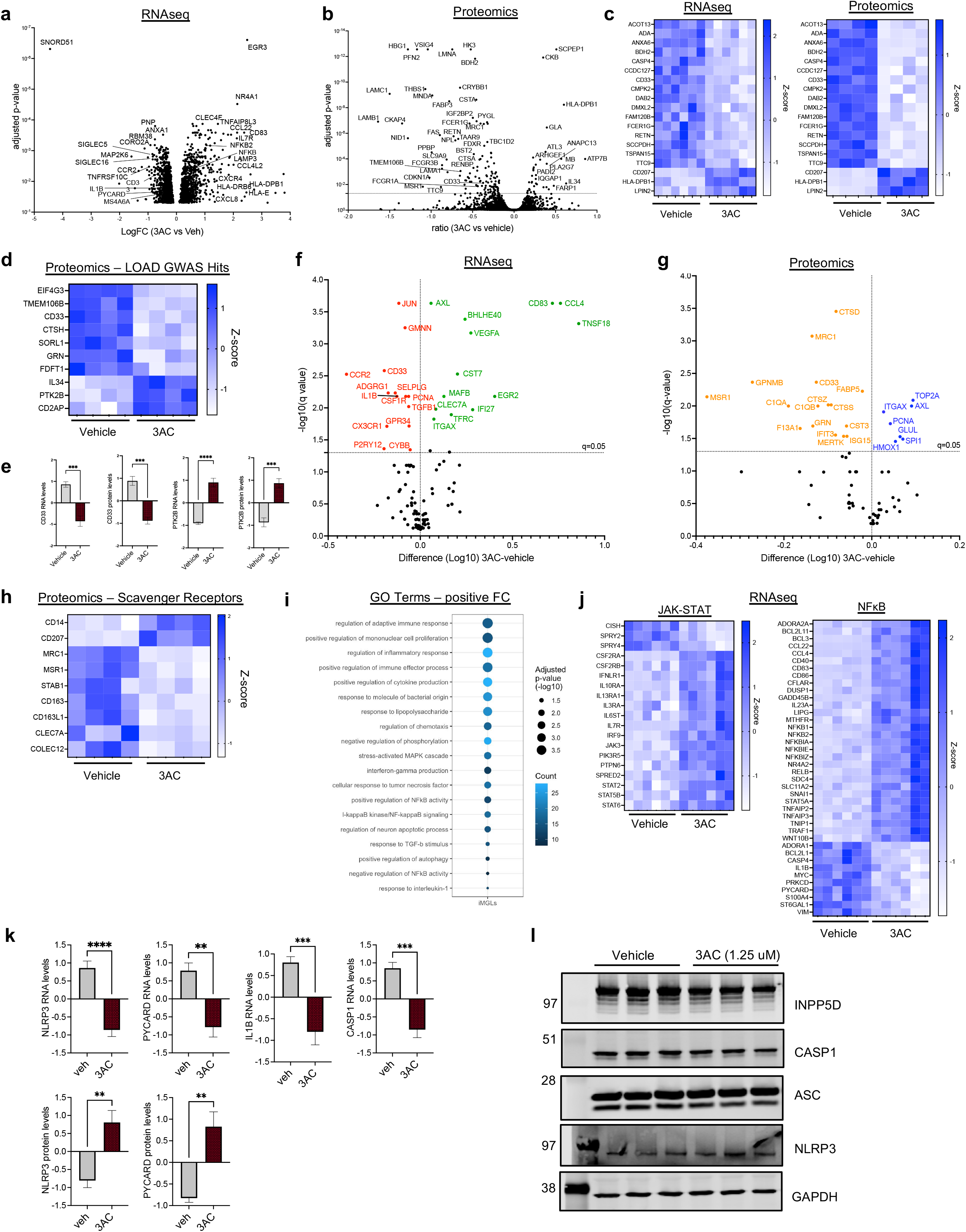
INPP5D inhibition induces changes in pathways associated with innate immunity in RNA and protein profiling. **a**. iMGLs were treated with vehicle (ethanol) or 3AC (1.25 uM) for 6 hours. Cells were then lysed, RNA purified, and RNAseq performed, n=6 per condition. Volcano plot of differentially expressed genes (DEGs) comparing vehicle and 3AC treatment conditions. 2,004 of 12,669 genes quantified were differentially expressed (Benjamini-Hochberg (BH), FDR<0.05) between vehicle and 3AC conditions. **b**. iMGLs were treated with vehicle (ethanol) or 3AC (1.25 uM) for 6 hours. Cells were then lysed in urea and mass spectrometry performed. n=4 per condition. Volcano plot of differentially expressed proteins (DEPs); 159 of 7,866 proteins were differentially expressed with 3AC treatment (BH FDR<0.05). **c**. Heat map of expression of DEGs that showed concordant differential expression at the RNA and protein level. **d**. Heatmap of relative expression of DEPs encoded by LOAD GWAS candidate genes^10^ between vehicle and 3AC treated iMGLs. **e**. Relative RNA and protein levels of CD33 and PTK2B between vehicle and 3AC treated microglia, as quantified by RNA sequencing and MS. Mean +/-SEM. Welch’s t-test. **f-g**. A variety of microglial subtypes previously have been defined through single-cell sequencing, with specific subtypes implicated to be altered in AD brain and model systems^34-38^. Volcano plots comparing 3AC vs vehicle treatment for genes defining these subtypes within the transcriptomic and proteomic data sets; significance determined by BH FDR<0.05. **h**. Heatmap of relative protein-levels of scavenger receptors in vehicle and 3AC treated iMGLs. **i**. Enriched gene ontology (GO) terms of biological processes for the differentially expressed genes that are elevated with 3AC treatment (geneontology.com^39^). **j**. Heatmap of relative expression between vehicle and 3AC-treatment of DEGs associated with NFΚB signaling and JAK-STAT signaling. **k**. Relative RNA and protein levels of inflammasome related components—NLRP3, PYCARD, IL1B, CASP1—in vehicle and 3AC-treated iMGLs, graphed by z-score. Mean +/-SEM. Welch’s t-test. **l**. Western blot of iMGL treated with either vehicle (ethanol) or 3AC (1.25 uM) for 6 hours showing protein expression of INPP5D and inflammasome related proteins: CASP1, ASC, and NLRP3, as well as GAPDH. Cell lines used for experiments are detailed in **Supplement Table 3**. For e and k: **p < 0.01, *** p < 0.001, **** p < 0.0001

A variety of microglial subtypes have been defined through single-cell sequencing, with specific subtypes implicated in AD brain and model systems (for example, homeostatic, disease-associated microglia “dam”, microglial neurodegenerative phenotype “mgnd”)^45–49^. We examined expression of genes defining these subtypes within the transcriptomic and proteomic data sets (**Fig. 2f-g**). We observe a protein-level decrease of lysosomal proteins CTSD and CTSS and of complement proteins C1QA and C1QB in response to INPP5D inhibition. Intriguingly we also observed a strong decrease in cell surface proteins such as MRC1 and MSR1 with 3AC treatment. Both MRC1 and MSR1 are categorized as scavenger receptors and several other scavenger receptors also showed a protein-level decrease with 3AC treatment (**Fig. 2h**). While expression of several of these microglia-subtype marker genes were altered with INPP5D inhibition, we did not observe a clear shift (that was consistent across several genes) from one defined subtype to another with INPP5D inhibition. We next used GeneOntology analyses (geneontology.org^50^) to characterize biological processes pathways affected by INPP5D inhibition. Pathways upregulated with 3AC included regulation of autophagy, regulation of cytokine production, and NFkB signaling (**Fig. 2i-j**). As NFkB signaling plays a critical role in mediating inflammatory responses, we examined the RNA and protein level changes of genes central to this pathway. Interestingly, key components of inflammasome signaling such as PYCARD (ASC), CASP1, IL1B, and NLRP3 were differentially expressed at the RNA and/or protein level (**Fig. 2k**). Western blotting confirmed the expression of inflammasome components CASP1, ASC, and NLRP3 in iMGLs at baseline and with 3AC treatment (**Fig 2l**). These changes following INPP5D inhibition implicate changes in innate signaling pathways and suggest a reduction in INPP5D activity may lead to inflammasome dysregulation.

### INPP5D inhibition results in inflammasome activation

To test whether INPP5D inhibition impacts profiles of secreted cytokines, iMGLs were treated with 3AC (1.25 uM) and cytokine response assayed. Conditioned media following a six-hour treatment were run on a panel for 10 proinflammatory cytokines (MesoScale Discoveries, IFN-γ, IL-1ß, IL-2, IL-4, IL-6, IL-8, IL-10, IL-12p70, IL-13, and TNF-α). Levels of cytokines were normalized to vehicle-treated cells and benchmarked against effects of treatment with lipopolysaccharide (LPS) (**Fig. 3a**). LPS-treated iMGLs resulted in dramatic (up to 300-fold) increases in TNF, IL-6, IL-8, IL-10, and IL-13. With 3AC-treatment, several cytokines were elevated, but IL-1ß showed the highest fold change in secretion that was consistently observed across multiple iMGL differentiations and multiple genetic backgrounds (**Fig. 3a, Extended Data Fig. 2**).

**Figure 3:**
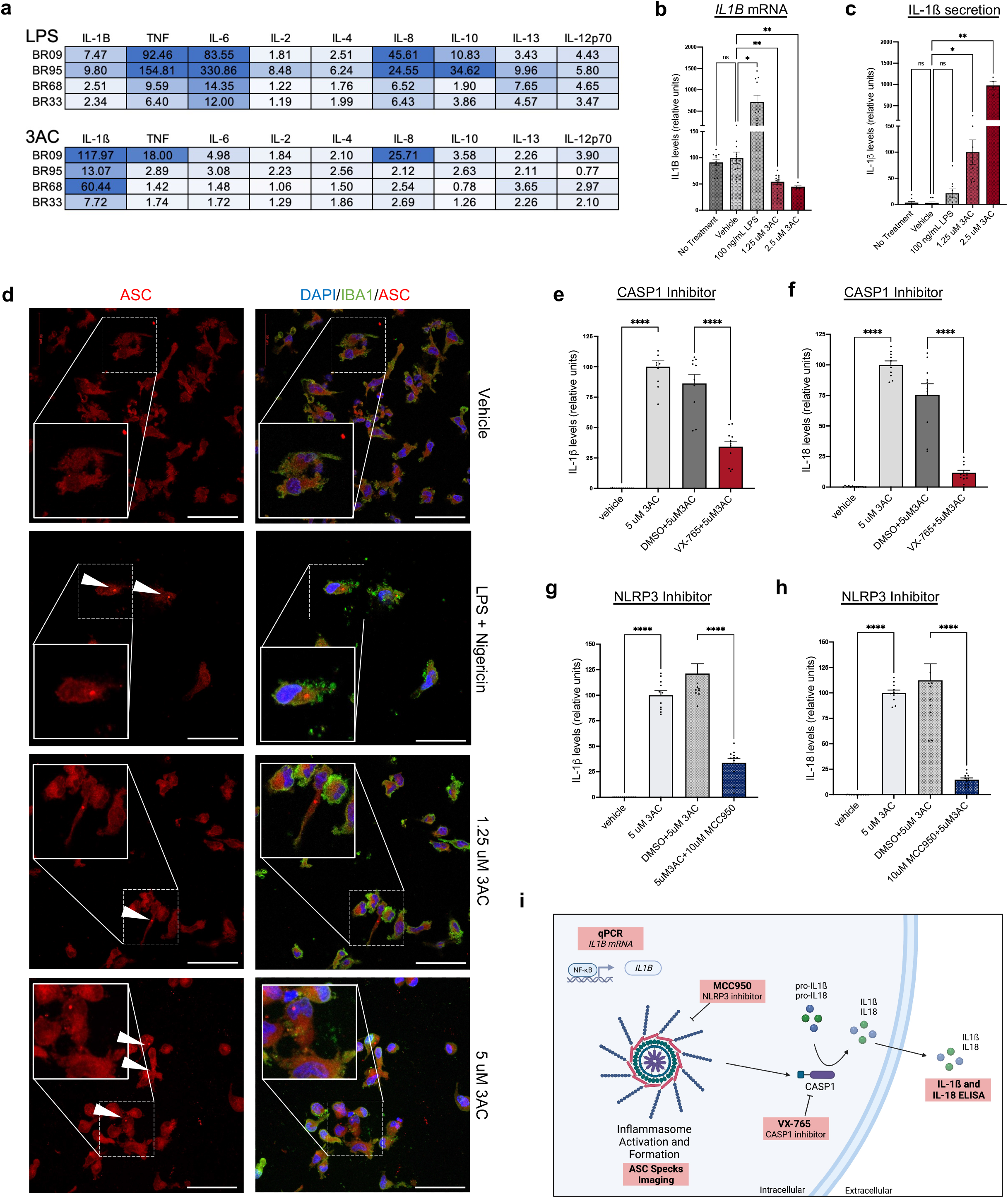
Acute INPP5D inhibition results in inflammasome activation and an increase in the secretion of IL-1ß and IL-18 in iMGL cultures. **a**. Fold change of secreted cytokines from iMGLs treated with LPS (100 ng/mL) or 3AC (1.25 uM) for 6 hours across iMGLs derived from four iPSC lines, assayed on a multiplex pro-inflammatory MSD ELISA assay. Fold change was calculated compared to vehicle (ethanol) treated cells. **b**. Levels of *IL1B* following 6-hour treatment with vehicle (ethanol), LPS (100 ng/mL), or 3AC (1.25 uM, 2.5 uM) treatment as measured by quantitative real-time PCR (qPCR). Data are normalized to *GAPDH* expression then values were normalized to vehicle treatment within each experiment. 3 differentiations, n=3-4 per condition. Mean +/-SEM. One-way ANOVA with Dunnett’s T3 multiple comparisons test. **c**. Secreted levels of IL-1ß measured in the conditioned media for the same experiments as (b) following 6-hour treatment with vehicle (ethanol), LPS (100 ng/mL), or 3AC (1.25 uM, 2.5 uM) as measured by ELISA. IL-1ß levels were normalized to 1.25 uM 3AC treated conditions within each experiment. 3 differentiations, n=3-4 per condition. Mean +/-SEM. One-way ANOVA with Dunnett’s T3 multiple comparisons test. **d**. iMGLs treated with either vehicle (ethanol), primed with LPS (100 ng/mL) for 3 hours and then treated with nigericin (10 uM) for 1 hour, or treated with 3AC (1.25 uM or 5 uM) for 2 hours. Cells were immunostained for ASC and IBA1. DNA is stained with DAPI. Imaged using confocal microscopy. Scale bars = 100 um. **e-f**. Treatment of iMGLs with either vehicle (ethanol) or 3AC (5 uM) with either VX-765 (25 uM) or its vehicle (DMSO). Levels of secreted IL-1ß and IL-18 were measured using an MSD dual IL-1ß and IL-18 ELISA following treatments and normalized to 3AC-treated samples in each experiment. Mean +/-SEM. One-way ANOVA with Sidak’s multiple comparison test. 3 differentiations, n = 10 per condition. **g-h**. Treatment of iMGLs with either vehicle (ethanol) or 3AC (5 uM) with either MCC950 (10 uM) or its vehicle (DMSO). Levels of secreted IL-1ß and IL-18 in the conditioned media were measured using an MSD dual IL-1ß and IL-18 ELISA and normalized to 3AC-treated samples in each experiment. Mean +/-SEM. One-way ANOVA with Sidak’s multiple comparison test. 4 differentiations, n=5-9 per condition. **i**. A summary figure of experiments performed to interrogate inflammasome activation, created using BioRender.com. For b-c, e-h: ns = not significant, *p < 0.05, ** p < 0.01, *** p < 0.001, **** p < 0.0001 Cell lines used for experiments are detailed in **Supplement Table 3**.

We next measured *IL1B* mRNA levels following treatment with 3AC (1.25 uM, 6 hours) to determine whether INPP5D activity affects *IL1B* transcription. iMGLs were treated with LPS (100 ng/mL, 6 hours) in parallel as a positive control for upregulation of *IL1B* mRNA, and *IL1B* mRNA measured by quantitative RT-PCR (qPCR). Surprisingly, *IL1B* mRNA levels did not increase following 3AC treatment. Rather, 3AC treatment induced a *reduction* in *IL1B* mRNA while increasing IL-1ß secretion (**Fig. 3b-c**), signifying that INPP5D inhibition induces IL-1ß post-translational processing and secretion while inducing a negative feedback loop. The inflammasome, an innate immune sensing multimeric oligomeric complex, contributes to the maturation of IL-1ß and IL-18 primarily through caspase-1 cleavage and activation. As IL-18 is also cleaved and activated in a similar manner as IL-1ß, we measured the secretion of IL-18 in the conditioned media of 3AC-treated iMGLs by ELISA (**Extended Data Fig. 2b-c**). Like IL-1ß, INPP5D inhibition induced secretion of IL-18 across iMGLs of different genetic backgrounds (**Extended Data Fig. 2c**). 3AC treatment of other iPSC-derived cell types that do not express INPP5D (iNs and iAs) did not yield an increase in IL-1ß or IL-18 secretion, supporting the specificity of the effect of 3AC on INPP5D at the low concentration used here (**Extended Data Fig. 3**).

Following its formation, the inflammasome cleaves pro-caspase-1 to generate the active form of caspase-1. Western blotting of iMGL lysates revealed an increase in cleaved caspase-1 with acute (2-3 hour) INPP5D inhibition with 3AC (**Extended Data Fig. 4a**). We reasoned that if active caspase-1 is involved in the INPP5D-inhibition-induced increase of IL-1ß and IL-18 secretion, then co-treatment of iMGLs with both 3AC and a caspase-1 inhibitor would result in decreased IL-1ß and IL-18 secretion relative to 3AC treatment alone. iMGLs were treated with either vehicle (ethanol) or 3AC (5 uM) along with either vehicle (DMSO) or caspase-1 inhibitor VX-765 (25 uM) for 6 hours. Conditioned media were collected and levels of IL-1ß and IL-18 secretion measured by ELISA. Indeed, co-treatment with VX-765 prevented the INPP5D-inhibitor induced elevation in IL-1ß and IL-18 (**Fig. 3e-f**). Treatment with a second caspase-1 inhibitor, Ac-YVAD-cmk, also prevented the 3AC-induced elevation IL-1ß and IL-18 secretion (**Extended Data Fig. 4b-c**). These results suggest that active caspase-1 is required for INPP5D-mediated effects on IL-1ß and IL-18 secretion.

**Figure 4.**
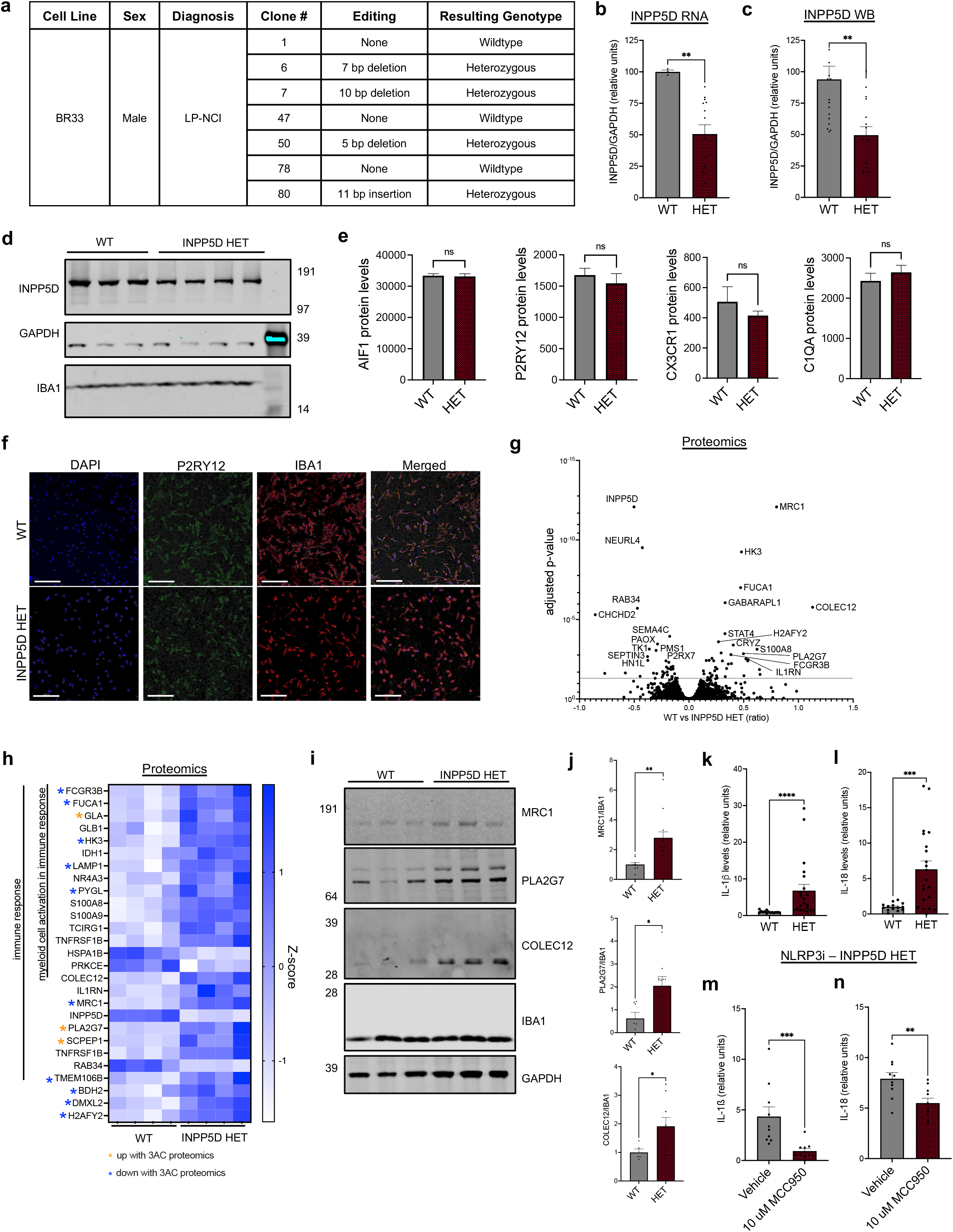
Chronic reduction in INPP5D levels results in an elevation of IL-1ß and IL-18. **a**. Table of CRISPR-Cas9 generated INPP5D heterozygous (HET) and monoclonally-selected wildtype (WT) lines. **b**. INPP5D RNA levels measured by qPCR, normalized to GAPDH levels. Mean +/-SEM. Mann-Whitney test. **c**. Western blot quantification of INPP5D protein levels in INPP5D WT and HET iMGLs normalized to GAPDH levels, Mean +/-SEM. Mann-Whitney test. **d**. Representative western blot of INPP5D, GAPDH, and IBA1 protein levels in INPP5D WT and HET iMGLs. **e**. INPP5D WT and HET iMGLs were lysed in urea and mass spectrometry performed, n=4 WT, n=4 HET. Relative protein levels of AIF1, P2RY12, CX3CR1, C1QA, as quantified by mass spectrometry. Mean +/-SEM. Unpaired t-test. **f**. Images of INPP5D WT and HET cells immunostained for IBA1 and P2RY12. DNA is stained with DAPI. Imaged using confocal microscopy. Scale bars = 200 um. **g**. Volcano plot of adjusted p-values showing 75 DEPs from the 7,868 quantified via TMT-MS (BH FDR<0.05). **h**. Heatmap of relative expression levels of proteins involved in immune signaling that are differentially up and down regulated (adj. p-value <0.05) in INPP5D WT vs HET iMGLs. Asterisks indicate proteins that also were differentially up or down regulated (q<0.05) with 3AC treatment (**Figure 2**). **i**. Representative western blot of MRC1, PLA2G7, COLEC12, IBA1, and GAPDH in INPP5D WT and HET iMGLs. **j**. Western blot quantification of MRC1, PLA2G7, and COLEC12 levels normalized to IBA1 levels. 3 differentiations. Mean +/-SEM; unpaired t-test. **k-l**. Secreted IL-1ß and IL-18 measured by ELISA of media collected from INPP5D HET and WT iMGLs. Media were concentrated 10-fold for detection and normalized to WT mean. n=14 wells WT, n=21 well HET. Mean +/-SEM. Mann-Whitney test. **m-n**. Secreted IL-1ß and IL-18 measured by ELISA of media collected from INPP5D HET iMGLs treated with either vehicle (DMSO) or MCC950 (10 uM) for 24 hours. Media were concentrated 10-fold for detection. n=10 wells vehicle, n=10 wells MCC950-treated. Mean +/-SEM. Mann-Whitney test. For b-c, e, j-n: ns = not significant, *p < 0.05, ** p < 0.01, *** p < 0.001, **** p< 0.0001

Immunostaining for the inflammasome adaptor protein, ASC (PYCARD), allows for visualization of inflammasome formation as the oligomerization of ASC within the inflammasome will form a “speck” within the cell^51^. As a positive control, iMGLs were first primed with LPS (100 ng/mL) for 3 hours and then treated with an inflammasome activator, nigericin (10 uM), for 1 hour prior to fixing and immunostaining for ASC. In parallel, iMGLs were treated with 3AC for 4 hours (1.25 uM, 5 uM) and fixed and immunostained for ASC. ASC specks of about 1 micron in size, consistent with what is reported in literature^51^, were detected in the iMGLs treated with LPS and nigericin and in the iMGLs treated with 3AC, while no ASC specks were observed in the vehicle treated conditions (**Fig. 3d**).

Next, we utilized an NLRP3 inhibitor, MCC950, to inhibit inflammasome formation^52^. MCC950 binds to NLRP3 and prevents a conformation change necessary for inflammasome formation, thus preventing inflammasome activation^53^. iMGLs were treated with either vehicle (ethanol) or 3AC (5 uM) along with either vehicle (DMSO) or MCC950 (10 uM) for 6 hours and then conditioned media were collected to measure levels of secreted IL-1ß and IL-18 (**Fig. 3g-h**). Inhibition of NLRP3-inflammasome formation prevented the effects of 3AC on IL-1ß and IL-18 secretion, signifying that inflammasome formation occurs downstream of INPP5D inhibition. Taken together, the visualization of ASC specks, CASP1 inhibitor treatments, and MCC950 treatment experiments support the hypothesis that INPP5D phosphatase activity suppresses inflammasome activation in iPSC-derived microglia (**Fig. 3i**).

### Chronic, genetically induced reduction of INPP5D induces protein-level changes in immune response proteins and increased secretion of IL-1ß and IL-18

The experiments outlined above propose that acute reduction in INPP5D phosphatase activity results in inflammasome activation. The reduction in INPP5D protein levels in AD brain likely results in chronic reduction of INPP5D activity which may have different outcomes compared to acute, efficient inhibition of INPP5D phosphatase activity. To interrogate the consequences of a chronic reduction of INPP5D activity, CRISPR-Cas9 was used to introduce loss-of-function mutations at the INPP5D locus in a single allele. The human iPSC line chosen was derived from a cognitively normal individual with minimal (lowest quartile within age 65+ cohort) neuritic plaque and tangle burden in their brain at death at age 90+years (BR33^30^). Heterozygous loss-of-function mutations (“HET”) were identified in several monoclonal lines, and wild type monoclonal lines also were isolated that were subjected to the same process. **Figure 4a** outlines the mutations introduced and clones isolated. In microglia derived from each of the heterozygous clones, INPP5D mRNA levels were reduced by approximately 50% (**Fig. 4b**) and protein levels were reduced by approximately 50% (**Fig. 4c-d**). Immunostaining for P2RY12 and IBA1 revealed that all cells expressed IBA1 and P2RY12, revealing that decreased INPP5D expression did not affect expression of these microglial markers (**Fig. 4f**).

We next examined the consequences of chronic loss-of-function of one allele of INPP5D on protein profiles. Expression of the microglial proteins AIF1, P2RY12, CX3CR1, and C1QA were not significantly changed between WT and HET iMGLs, demonstrating that one less functional copy of INPP5D did not alter the microglia fate of the cells (**Fig. 4e**) nor did it significantly affect viability (**Extended Data 5**). Differential protein analysis between WT and INPP5D HET iMGLs revealed 71 differentially expressed proteins—38 proteins were significantly increased in the INPP5D het iMGLs and 33 proteins were significantly decreased (**Fig. 4g, Extended Data Fig. 6a**). Downregulated proteins were not enriched in any GO terms, but include proteins involved in protein folding and stability such as TTC9, PDIA5, USOA1B and IRF2BPL, as well as proteins previously implicated in microglial dysfunction such as SEPTIN3 and P2RX7, the latter of which has been linked to inflammasome activation^54,55^. Proteins upregulated in INPP5D HET iMGLs include many falling under the GO term of “myeloid cell activation in immune response” such as PYGL, TCIRG1, GLA, FCGR3B, TNFRSF1B, HK3, S100A9, S100A8, GLB1, IDH1, FUCA1, LAMP1, NR4A3 (**Fig. 4h**). Through western blot analyses, we confirmed the significant upregulation of MRC1, PLA2G7 and COLEC12 in the INPP5D HET iMGLs (**Fig. 4i-j**). Upregulation of proteins such as IL1RN^56^, which are implicated as negative regulators of innate immune activation suggest there may be a negative feedback mechanism with a chronic reduction in INPP5D activity. Comparison of the DEPs between acute (vehicle vs 3AC) and chronic (WT vs INPP5D HET) reduction revealed 14 DEPs shared between the two analyses (**Extended Data Fig. 6b**). Intriguingly, 10 of the proteins that were downregulated with 3AC treatment were upregulated in the INPP5D het (BDH2, DMXL2, FCGR3B, FUCA1, H2AFY2, HK3, LAMP1, MRC1, PYGL, and TMEM106B), a number higher than would be expected by chance (Chi square test, p<1×10^−15^; **Extended Data Fig. 6b**). Thus, reduction in INPP5D activity may result in inflammasome activation and induction of a negative feedback mechanism that results in lowering of IL1B RNA levels (as in **Fig. 3b**). Despite this potential feedback mechanism, chronic decrease of INPP5D resulted in an elevation in extracellular secretion of IL-1ß and IL-18 (**Fig. 4k-l**). To determine whether this increase was the result of inflammasome activation, we treated INPP5D HET iMGLs with either vehicle (DMSO) or NLRP3 inhibitor, MCC950 (10 uM), for 24 hours. Following treatment, a significant decrease in IL-1ß and IL-18 secretion with MCC950 treatment was observed (**Fig. 4m-n**). Taken together, these data demonstrate that a chronic reduction of INPP5D results in both sub-lytic levels inflammasome activation and protein-level changes in immune response that are resulting from inflammasome activation.

## DISCUSSION

Our data suggest that understanding the interplay of signaling mechanisms between INPP5D activity and inflammasome regulation could be key to unraveling aspects of the early stages of microglial involvement in AD. By examining altered transcriptional and proteomic profiles resulting from decreased INPP5D activity, our study suggests three potentially overlapping avenues for linking INPP5D function to microglial activation and elevated risk for AD: 1) altered inflammasome activity within microglia; 2) immune sensing pathways that would affect microglia recognition and engulfment of Aß; and 3) as-of-yet undefined signaling mechanisms that affect the expression of other proteins linked to LOAD pathogenesis through GWAS.

We observed changes of several LOAD GWAS associated proteins in both the 3AC-treated and INPP5D HET models including CD33 and TMEM106B, while several other GWAS associated genes implicated in microglial biology were unchanged (for example, APOE and PLCG2). Both CD33 and TREM2 have been linked to INPP5D biology previously^15,57^. CD33 protein and RNA levels were reduced with acute inhibition of INPP5D activity, while TREM2 RNA levels were not affected (TREM2 was not detected in all samples at the protein level via MS). Altered levels of other LOAD GWAS related genes following decreased activity of INPP5D indicates that a number of these proteins likely interact functionally with one another.

Microglia recognition and engulfment of Aß peptides are implicated as important processes in the early stages of AD. Scavenger receptors are putative Aß receptors, important for recognition and uptake of Aß from the extracellular space by microglia. Proteomic and transcriptomic analysis of reduction of INPP5D activity acutely and chronically each revealed significant changes in the expression of scavenger receptors. With acute inhibition, protein levels of MRC1, SCARA1/MSR1, COLEC12, and CLEC7A were reduced. MSR1 is reported as an Aß receptor and decreased MSR1 expression by microglia leads to increased Aß accumulation^58^. This downregulation of scavenger receptors would lead to altered Aß clearance and an accumulation of Aß proteins. However, perhaps through the above-proposed feedback mechanism, chronic reduction of INPP5D levels resulted in an *elevation* in protein levels of the scavenger receptors MRC1, COLEC12, CD36 and CD163. CD36 binds to and enhances Aß clearance ^59^ and COLEC12 is upregulated in glia near Aß plaques in AD brain and in fAD transgenic mouse models^60^. Other than INPP5D itself, MRC1 was the protein most strongly affected by loss of one copy of INPP5D (**Fig. 4g**). In previous published studies, MRC1 (CD206) is upregulated in microglia following injury *in vivo* and following treatment with IL-4 *in vitro* ^61,62^. Thus, modulation of INPP5D activity impacts the profile of scavenger receptor present on the microglia, which in turn may affect both Aß clearance and phagocytosis of other extracellular factors.

Inflammasome activity has been linked to several diseases including AD, thus understanding its method of regulation is critical towards unraveling its role in a number of biological processes^63^. The mechanism of inflammasome regulation and activation appears to be complex and multifaceted, with several aspects remaining to be uncovered. Multiple therapeutic efforts are ongoing to develop and test pharmacologic inhibitors of the NLRP3 inflammasome (such as MCC950, used herein). Previous studies have shown that Aß can activate the inflammasome to increase IL-1ß secretion^45,64^, and that activation of the NLRP3 inflammasome affects plaque levels and contributes to spatial memory defects in APP/PS1 mice^4^. Recent studies linked the NLRP3 inflammasome to the development of tau pathology through phosphatase regulation by the NLRP3 inflammasome and through the worsening of tau pathology through IL-1ß signaling^5^. Here, we show activation of the inflammasome in the absence of external AD-related insults such as neurotoxic Aß or tau. Thus, the activation of the NLRP3 inflammasome following decreased INPP5D activity in microglia may contribute to risk for AD by altering the vulnerability of microglia to additional insults encountered in the brain. Additional studies in the coming years will be necessary to test this hypothesis.

Examining the differences between the vehicle versus 3AC-treatment and INPP5D WT versus HET comparisons provide a window into the mechanisms at play following an acute versus chronic decrease in INPP5D activity. With acute reduction of INPP5D activity, the inflammasome is activated with a strong induction of IL-1ß and IL-18 release (up to a 1000-fold increase with 2.5 uM 3AC, **Fig. 3c**). Extracellular levels of IL-1ß and IL-18 also are elevated with chronic reduction of INPP5D, although the effect is blunted in comparison to acute efficient inhibition (less than 10-fold increase, **Fig. 4k-l**). However, this increase is rescued with NLRP3 inhibition (**Fig. 4m-n**) demonstrating that a low level of inflammasome activation is contributing to this phenotype. With this chronic loss of INPP5D, we also observe changes that suggest a long-term feedback mechanism as the cells respond to inflammasome. This may be mediated in part by the observed upregulation of IL1RN protein, a negative regulator of IL-1R activity. Proteins shown to be rapidly reduced following acute INPP5D inhibition but then elevated with chronic INPP5D loss include MRC1, TMEM106B, PYGL, LAMP1, HK3, H2AFY2, FUCA1, FCGR3B, DMXL2, and BDH2 (**Extended Data Fig. 6b**).

Data presented herein provide clues to uncovering the signaling cascades that link INPP5D activity to inflammasome regulation. Through the integration of our findings in human microglia with other experimental systems reported in the literature, two candidate pathways emerge. One pathway implicated is through CLEC7A/Dectin-1 signaling and the second through PLA2G7. INPP5D is reported to affect C-Type Lectin Domain Containing 7A (CLEC7A) signaling through FcgammaR interactions in THP-1 and bone marrow cells^16,17^. In dendritic cells, CLEC7A is involved in a non-canonical inflammasome pathway to induce IL-1ß signaling^65^. Recent studies report that these CLEC7A/inflammasome pathways may be synergistic with the NLRP3 inflammasome^66^. Here, we observed a decrease in CLEC7A protein following INPP5D inhibition coupled to inflammasome activation, suggesting a linkage also exists between these pathways in human microglia (**Fig. 2h**). Secondly, Lipoprotein-associated Phospholipase A2 (PLA2G7) protein levels were rapidly elevated with INPP5D inhibition, and these elevated protein levels also were observed with chronic INPP5D reduction in HET microglia. This elevation was identified through unbiased protein profiling and was validated via western blotting (**Fig. 4i-j**). PLA2G7 functions by hydrolyzing glycerophospholipids to generate lysophosphatidylcholine (LysoPC), which has proinflammatory effects^67^. PLA2G7 has been primarily studied in its role in atherosclerosis, and an inhibitor of PLA2G7 activity, darapladib, has been proposed as a therapeutic target for atherosclerosis^68–70^. Intriguingly, a recent study showed that darapladib-mediated inhibition of PLA2G7 provides benefit through a downregulation of NLRP3 inflammasome activation in macrophages^71^. Our data show that decreased INPP5D activity elevates protein levels of PLA2G7 which in turn may induce inflammasome activation in human microglia as well. Future experimental efforts involve utilizing darapladib to investigate whether PLA2G7 inhibition rescues INPP5D inhibition-induced inflammasome activation.

We recognize limitations of the current study. These *in vitro* studies are reductionist in nature, with reduced complexity regarding cell type representation compared to the endogenous environment of the brain. Isolation of microglia away from other cell types can result in an artificial activation of the cells which in turn can affect expression profiles. Thus, it is important to interpret in vitro results in the context of findings from other systems. However, iPSC models also provide some advantages over *in vivo* systems including an increased cell yield, which allows for protein level analyses following pharmacological and genetic manipulations under well-controlled conditions. Here, integrating our findings with previously published findings from humans and mice aids in our ability to interpret the results. Through the work that is presented in this study, we link INPP5D activity to immune processes in microglia and identify INPP5D as a novel regulator of NLRP3 inflammasome activity. As NLRP3 inflammasome activity has been linked to many diseases and disorders, the identification of INPP5D as a member of this pathway would have therapeutic impact in other disease fields beyond Alzheimer’s disease. In the coming years, the connection of INPP5D to the inflammasome pathway can be deeply studied using these and other experimental systems to further define members of this signaling cascade.

## METHODS

### Brain Extract Preparation

All work was performed following IRB review and approval through Partners/BWH IRB (2016P000867). Human brain material was obtained from 1) the neuropathology core facility at Massachusetts General Hospital, 2) Rush University Medical Center, and 3) Albany Medical Center. ROS and MAP studies were approved by an Institutional Review Board of Rush University Medical Center. All participants signed an informed consent, an Anatomical Gift Act, and a repository consent to allow their data and biospecimens to be repurposed. IPSC lines were utilized following IRB review and approval through MGB/BWH IRB (#2015P001676). Tris Buffered Saline (TBS: 20 mM Tris-HCl, 150 mM NaCl, pH 7.4) brain extracts were prepared using the method previously described^30^. Briefly, the tissue was dissected to isolate the gray matter, which was homogenized in ice-cold TBS at a ratio of 1:4 tissue weight to buffer volume within a Dounce homogenizer. This suspension was then subject to ultra-centrifugation at 1.75 × 10^5^ x *g* in a TL100 centrifuge (Beckman Coulter) to pellet cellular debris. Urea brain extracts were prepared in 8M urea and prepared as per a previous study^23^. Experimenter was blinded to diagnosis for Western blotting quantification, wherein wells were excluded that showed “pinched” bands or abnormally high background. Those samples were re-run and quantified. Following quantification, outliers were eliminated with ROUT Q=10%, and then levels of INPP5D and GAPDH normalized to internal controls across blots.

### Immunostaining of human brain tissue

Tissue was dissected from the temporal cortex of post-mortem human brain tissue and drop-fixed in 10% formalin for 24 hours then buffer exchanged into 30% sucrose for cryopreservation. Tissue was embedded into Tissue-Tek OCT Compound (Sakura Finetek) and frozen for cryosectioning. Tissue was sectioned into 25-micron sections using a cryostat. Antigen retrieval was performed in citrate buffer prior to blocking with 3% BSA and 0.1% triton in PBS. Primary antibodies were prepared in 1% BSA in PBS (SHIP1, 1:100 dilution, Cell Signaling Technologies; IBA1, 1:500 dilution, abcam) and sections were incubated overnight in this solution at 4°C. Washes were performed with PBS and incubated in secondaries (Cy2 Donkey anti-Rabbit or Cy3 Donkey anti-Goat, 1:2000 dilution, Jackson Immunoresearch) diluted in PBS for 1 hour at room temperature. After washing, sections were incubated in Sudan black in 70% ethanol for 10 minutes. Sections were washed in PBS and mounted onto glass slides using Vectashield with DAPI for confocal microscopy using a Zeiss LSM710 confocal microscope and acquired using ZEN black software.

### Immunoblotting

iMGL samples were harvested with NP40 buffer (1% Nondeit 40, 0.15 mM NaCl, 5 mM Tris, pH 7.4, 1 mM EDTA) supplemented with cOmplete protease inhibitor cocktail (Roche) and PhoSTOP phosphatase inhibitors (Roche). Samples were spun down at 15,000 x *g* for 15 minutes at 4°C on a tabletop centrifuge and then the supernatant was collected for protein quantification using the Pierce BCA protein assay kit. Samples were prepared using 4X Protein Sample Loading Buffer (Licor) as per manufacturer instructions and then run on NuPAGE 4-12% Bis-Tris mini or midi protein gels with either 1X MOPS or 1X MES buffer. Western blot transfer was performed on ice using Tris Glycine transfer buffer onto nitrocellulose membrane and then blocked for 1 hour at room temperature with Intercept Blocking Buffer (Licor). Membranes were incubated with primary antibodies diluted in blocking buffer over night at 4°C (SHIP1, 1:500 dilution, Cell Signaling Technologies; IBA1, 1:500 dilution, abcam; GAPDH, 1:10,000 dilution, Millipore; PAFAH/PLA2G7, 1:500 dilution, Proteintech; MRC1; 1:1000 dilution, abcam; COLEC12, 1:500 dilution, R&D Systems; CASP1, 1:1000 dilution, Cell Signaling Technologies; ASC, 1;1000 dilution, Cell Signaling Technologies; NLRP3, 1:1000 dilution, Cell Signaling Technologies). Washes were performed with TBS buffer with 0.01% Tween-20 (TBS-T). Blots were incubated with secondary antibodies diluted in TBS-T buffer (800CW Donkey anti-Rabbit, 800CW Donkey anti-Goat, 680RD Donkey anti-Mouse, 1:10,000 dilution, Licor Biosciences) for 1 hour at room temperature. Washes were performed again in TBS-T before final storage in TBS for imaging using an LICOR Odyssey CLx imager. Bands were quantified using Image Studio Lite and signals normalized to GAPDH or IBA1 levels. For western blot analysis for Urea or TBS brain extracts, experimenters performing the western blots were blind to AD diagnosis of patient samples and ROUT Q=10% was used to eliminate outliers

### Generation of iMGLs from iPSCs

iPSCs were maintained on growth-factor reduced matrigel (Corning) and fed with StemFlex media (ThemoFisher Scientific) and undergo monthly mycoplasma testing to ensure cultures are mycoplasma-free with the LookOut Mycoplasma PCR detection kit (Millipore Sigma). For the differentiation of iPSC-derived microglia-like cells, iPSCs were differentiated using a previously established protocol^26,27^ that was further optimized, as described here. HPCs were generated from iPSCs using the StemDiff Hematopoietic Kit (StemCell Technologies). Cells were replated using ReLeSR (StemCell Technologies) at day 12 at 100,000 cells per 35 mm well (1 well of a 6 well plate). Cells were plated in 2 mL of iMGL media per well: DMEM/ F12, 2X insulin-transferrin-selenite, 2X B27, 0.5X N2, 1X glutamax, 1X non-essential amino acids, 400μM monothioglycerol, 5 μg/mL insulin. Microglia media is supplemented with fresh cytokines before each use: 100 ng/mL IL-34 (Peprotech), 50 ng/mL TGFβ1 (Millitenyi Biotech), and 25 ng/mL M-CSF (ThermoFisher Scientific). On days 14, 16, 18, 20, and 22, each well was supplemented with 1 mL of iMGL media with freshly added cytokines. On day 24, all media is removed from the wells and the cells are dissociated by incubating with 5 minutes of PBS at room temperature. Both the media and the PBS is centrifuged at 300 x *g* for 5 minutes to pellet adherent and non-adherent cells. Cells are combined and replated at 100,000 cells per 15.6 mm well (1 well of a 24 well plate) in 1 mL of a 1:1 mixture of old media and fresh iMGL media with tricytokine cocktail. On days 26, 28, 30, 32, 34, 36, each well is supplemented with 0.5 mL of iMGL media with freshly added 3 cytokines. On day 37, all but 0.5 mL of media was removed from each well. Media was centrifuged for 5 minutes at 300 x *g* to pellet non-adherent cells. Cells are resuspended in iMGL media supplemented with 100 ng/mL IL-34, 50 ng/mL TGFβ1, 25 ng/mL M-CSF, 100 ng/mL CD200 (Novoprotein) and 100 ng/mL CX3CL1 (Peprotech) for a total of 1 mL per well. On day 39, cells are fed with microglia media with five cytokine cocktail (0.5 mL per well). After day 40, cells were ready for experimentation.

### Immunocytochemistry of iMGLs

iMGLs were plated on either glass pre-coated coverslips or 96 well plates at a density of 50,000 cells per coverslip or 16,000 cells per well on day 24 with dissociation as explained above. Cells were fixed with 4% paraformaldehyde and then blocked for 1 hour at room temperature with blocking buffer (2% donkey serum, 0.1% triton, in PBS). Primary antibodies were prepared in blocking buffer (SHIP1, 1:100 dilution, Cell Signaling Technologies; IBA1, 1:500 dilution, abcam; ASC, 1:500 dilution, Cell Signaling Technologies; P2RY12, 1:500 dilution, gift from Oleg Butovsky’s lab) and fixed cells were incubated overnight in primary antibody at 4°C. Following gentle washes, secondaries diluted in PBS (Cy2 Donkey anti-Rabbit or Cy3 Donkey anti-Goat, 1:2000 dilution, Jackson Immunoresearch) and added for 1 hour at room temperature. For cells plated in wells, DAPI was added for a 10-minute incubation before final storage in PBS. Coverslips were mounted using Vectashield mounting medium with DAPI. Images were taken using a Zeiss LSM710 confocal microscope and acquired using ZEN black software. Blinding was applied to the examination of ASC spec formation in iMGLs with positive controls, negative controls, and 3AC treatment.

### Dissociation for Single Nuclear RNA Sequencing and library generation

All media was removed from the wells and the cells are dissociated by incubating with 5 minutes of PBS at room temperature. Both the media and the PBS are centrifuged for 5 minutes at 300 x *g* and the pellets were combined and washed once in ice-cold PBS with 0.04% BSA. The suspension was centrifuged for 5 minutes at 500 x *g* to pellet cells that were then lysed on ice for 15 minutes with lysis buffer: 10 mM Tris, 0.49% CHAPS, 0.1% BSA, 21 mM magnesium chloride, 1 mM calcium chloride, 146 mM sodium chloride. The nuclei were pelleted through centrifugation 5 minutes at 500 x *g* and resuspended in PBS + 1% BSA for counting prior to loading into the 10X following manufacturer’s instructions.

Single nucleus suspensions at 1k cells/uL were used to generate gel emulsion bead suspensions using the 10xGenomics Chromium controller and NextGEM 3’ reagents v3.1 (10X Genomics, San Francisco, CA) with a targeted recovery of 10000 cells. Single cell libraries were generated following the “Chromium NextGEM” protocol (CG000204 Rev D). A TapeStation 4200 was used to assay library quantity and size distribution. Libraries were sequenced at the New York Genome Center using a NovaSeq at a depth of 4×10^8^ reads per sample (4×10^4^ reads per cell). Fastq files were processed using the 10xGenomics CellRanger pipeline and standard CellRanger outputs were visualized using the Loupe Browser.

### Single Cell Analysis

The 10x Genomics CellRanger pipeline “filtered_feature_bc_matrix” output from two iPSC-derived microglia sample runs (Batches) were merged for downstream single nucleus RNA-seq data analysis using the Seurat v4.0.3 package for R. The data object was filtered by keeping only genes detected in at least 10 nuclei, and nuclei were filtered to only include those with detection of at least 100 genes. The percentages of mitochondrial genes and ribosomal protein-related genes were calculated, after which mitochondrial related genes and sex chromosome related genes (XIST, UTY) were removed. The data object was further filtered, keeping only nuclei with less than 5% mitochondrial related genes, with more than 1,000 and less than 25,000 unique mapped transcripts (Unique Molecular Identifiers – UMIs).

To assess cell type similarity of microglia, post-mortem human nuclei from individuals without Alzheimer’s Disease diagnoses were used, from the Cain et al. publication^33^. Glutamatergic neurons, astrocytes and microglia from this post-mortem data set were reprocessed by removing mitochondrial related genes and sex chromosome related genes (XIST, UTY), and these nuclei were then merged with the iMGL data object.

For normalization and integration of this merged data object, the Seurat function “SCTransform” was used to identify 3000 variable genes in all nuclei and regress out UMI counts. 25 principal components were used to resolve features in the merged data object and the Harmony function “RunHarmony” was used to integrate nuclei anchored by cell types. 25 Harmony-derived dimensions and 30 k-nearest neighbors were used to embed nuclei into a Uniform Manifold Approximation and Projection (UMAP) space for visualization. In parallel, these Harmony dimensions were used to identify cell clusters using the standard Louvain community detection algorithm implemented in Seurat. Finally, the Seurat function “FindAllMarkers” was used to find gene markers for cell clusters among the 3000 variable genes identified by SCTransform.

### iMGL Treatments and Cytokine measurements

3AC (Echelon Biosciences) was reconstituted in 100% ethanol as per manufacturer instructions. 3AC (1.25 uM, 2.5 uM, or 5 uM) treatment was performed for 6 hours. LPS, Nigericin, VX-765 (Invivogen), Ac-YVAD-cmk (Invivogen), and MCC950 (Invivogen) were reconstituted as per manufacturer instructions. iMGLs were treated with LPS (100 ng/mL) for 6 hours and VX-765 (25 uM) and MCC950 (10 uM) were treated with 3AC (5 uM) for 6 hours prior to harvesting. For Ac-YVAD-cmk treatment, iMGLs were pre-treated first with Ac-YVAD-cmk (20, 40, 80 uM) prior to 3AC (5 uM) treatment for 6 hours and harvesting. For ASC immunostaining, cells were treated with LPS (100 ng/mL) for 3 hours and then Nigericin (10 uM) for 1 hour prior to fixation with 4% paraformaldehyde. Conditioned media was collected from iMGLs following treatment. Conditioned media was assayed for cytokine levels using Meso Scale Discovery V-PLEX proinflammatory panel or the U-PLEX dual IL-1ß and IL-18 assay. For INPP5D WT and HET conditioned media, the media was concentrated at room temp using a Savant SpeedVac SPD1030 Vacuum Concentrator for 2 hours to concentrate the media 10-fold prior to loading onto the dual IL-1ß and IL-18 assay.

### Bulk RNA sequencing

For INPP5D acute inhibition, samples include iMGLs from 2 genetic backgrounds (BR01, BR33), treated with 3AC (1.25 uM 3AC, 6 hours) or vehicle (ethanol). INPP5D CRISPR reduction was in BR33 background. For all samples 500ngs of total RNA input was sequenced through the Genewiz polyA selection, HiSeq 2×150 single index sequencing platform. RNAseq reads were quality tested using fastqc, quality trimmed then quantified using the Kallisto pseudoalignment quantification program (v0.43.1)^72^ running 50 bootstraps against a Kallisto index generated from GRCh38. Kallisto quantified samples were analyzed using the “Sleuth” package (v0.30.0) in R Studio (v3.6.1 of R; v1.2.5019 of R Studio)^73^. Expression values were exported from the Sleuth object as normalized TPM values. The final RNAseq master expression matrix has 16 samples and quantifies expression of 43,122 genes. To identify differentially expressed genes the above matrix was filtered to remove low expressers (greater that 5TPM in at least 2 samples), leaving 13,532 quantified genes. Conditions were compared by linear modeling with empirical Bayesian analysis using the “limma” package^74^.

### TMT proteomics and data analysis

#### Sample Processing

Media was removed from the wells and iMGLs were dissociated by incubating with 5 minutes of PBS at room temperature. Cells were pelleted with a 500 x *g*, 5 minute spin at room temperature, and flash frozen in liquid nitrogen for transport. Each cell pellet was individually homogenized in 300 uL of urea lysis buffer (8M urea, 100 mM NaHPO4, pH 8.5), including 5 uL (100x stock) HALT protease and phosphatase inhibitor cocktail (Pierce). All homogenization was performed using a Bullet Blender (Next Advance) according to manufacturer protocols. Briefly, each tissue piece was added to Urea lysis buffer in a 1.5 mL Rino tube (Next Advance) harboring 750 mg stainless steel beads (0.9-2 mm in diameter) and blended twice for 5 minute intervals in the cold room (4°C). Protein supernatants were transferred to 1.5 mL Eppendorf tubes and sonicated (Sonic Dismembrator, Fisher Scientific) 3 times for 5 s with 15 s intervals of rest at 30% amplitude to disrupt nucleic acids and subsequently vortexed. Protein concentration was determined by the bicinchoninic acid (BCA) method, and samples were frozen in aliquots at −80°C. Protein homogenates (50ug) treated with 1 mM dithiothreitol (DTT) at 25°C for 30 minutes, followed by 5 mM iodoacetimide (IAA) at 25°C for 30 minutes in the dark. Protein mixture was digested overnight with 1:100 (w/w) lysyl endopeptidase (Wako) at room temperature. The samples were then diluted with 50 mM NH4HCO3 to a final concentration of less than 2M urea and then and further digested overnight with 1:50 (w/w) trypsin (Promega) at 25°C. Resulting peptides were desalted with a Sep-Pak C18 column (Waters) and dried under vacuum.

#### Tandem Mass Tag (TMT) Labeling

Peptides were reconstituted in 100ul of 100mM triethyl ammonium bicarbonate (TEAB) and labeling performed as previously described^75,76^ using TMTPro isobaric tags (Thermofisher Scientific, A44520). Briefly, the TMT labeling reagents were equilibrated to room temperature, and anhydrous ACN (200 μL) was added to each reagent channel. Each channel was gently vortexed for 5 min, and then 20 μL from each TMT channel was transferred to the peptide solutions and allowed to incubate for 1 h at room temperature. The reaction was quenched with 5% (vol/vol) hydroxylamine (5 μl) (Pierce). All 16 channels were then combined and dried by SpeedVac (LabConco) to approximately 100 μL and diluted with 1 mL of 0.1% (vol/vol) TFA, then acidified to a final concentration of 1% (vol/vol) FA and 0.1% (vol/vol) TFA. Peptides were desalted with a 60 mg HLB plate (Waters). The eluates were then dried to completeness.

#### High pH Fractionation

High pH fractionation was performed essentially as described^77^ with slight modification. Dried samples were re-suspended in high pH loading buffer (0.07% vol/vol NH4OH, 0.045% vol/vol FA, 2% vol/vol ACN) and loaded onto a Water’s BEH (2.1mm x 150 mm with 1.7 µm beads). An Thermo Vanquish UPLC system was used to carry out the fractionation. Solvent A consisted of 0.0175% (vol/vol) NH4OH, 0.01125% (vol/vol) FA, and 2% (vol/vol) ACN; solvent B consisted of 0.0175% (vol/vol) NH4OH, 0.01125% (vol/vol) FA, and 90% (vol/vol) ACN. The sample elution was performed over a 25 min gradient with a flow rate of 0.6 mL/min with a gradient from 0 to 50% B. A total of 96 individual equal volume fractions were collected across the gradient and dried to completeness using a vacuum centrifugation.

#### Liquid Chromatography Tandem Mass Spectrometry

All samples were analyzed on the Evosep One system using an in-house packed 15 cm, 75 μm i.d. capillary column with 1.9 μm Reprosil-Pur C18 beads (Dr. Maisch, Ammerbuch, Germany) using the pre-programmed 21 min gradient (60 samples per day) essentially as described^78^. Mass spectrometry was performed with a high-field asymmetric waveform ion mobility spectrometry (FAIMS) Pro equipped Orbitrap Eclipse (Thermo) in positive ion mode using data-dependent acquisition with 2 second top speed cycles. Each cycle consisted of one full MS scan followed by as many MS/MS events that could fit within the given 2 second cycle time limit. MS scans were collected at a resolution of 120,000 (410-1600 m/z range, 4×10^5 AGC, 50 ms maximum ion injection time, FAIMS compensation voltage of - 45). All higher energy collision-induced dissociation (HCD) MS/MS spectra were acquired at a resolution of 30,000 (0.7 m/z isolation width, 35% collision energy, 1.25×10^5 AGC target, 54 ms maximum ion time, TurboTMT on). Dynamic exclusion was set to exclude previously sequenced peaks for 20 seconds within a 10-ppm isolation window. Data Processing Protocol

All raw files were searched using Thermo’s Proteome Discoverer suite (version 2.4.1.15) with Sequest HT. The spectra were searched against a human uniprot database downloaded August 2020 (86395 target sequences). Search parameters included 10ppm precursor mass window, 0.05 Da product mass window, dynamic modifications methione (+15.995 Da), deamidated asparagine and glutamine (+0.984 Da), phosphorylated serine, threonine, and tyrosine (+79.966 Da), and static modifications for carbamidomethyl cysteines (+57.021 Da) and N-terminal and Lysine-tagged TMT (+304.207 Da). Percolator was used filter PSMs to 0.1%. Peptides were group using strict parsimony and only razor and unique peptides were used for protein level quantitation. Reporter ions were quantified from MS2 scans using an integration tolerance of 20 ppm with the most confident centroid setting. Only unique and razor (i.e., parsimonious) peptides were considered for quantification.

Proteomic data were filtered to remove any proteins with missing. The ComBat algorithm was used to remove variance induced by differentiation round. Differentially Expressed Proteins were identified by linear modeling with empirical Bayesian analysis using the “DEP” package^79^ in R.

### Real-Time Quantitative PCR

RNA was harvested from iMGLs and purified using kit instructions (PureLink RNA Mini Kit, Invitrogen). cDNA was generated from RNA using SuperScript II Reverse Transcriptase. Assay was performed using Power SYBR™ Green PCR Master Mix on an Applied Biosciences Vii7a Real-time PCR machine. Samples were assayed with 3 technical replicates and analyzed using the DDC_T_ method and expression was normalized to GAPDH expression. All primer sequences are detailed in **Supplementary Table 4**.

### CRISPR targeting to generate INPP5D heterozygote cells

Guide RNAs were designed using the Broad Institute CRISPick sgRNA design tool^80,81^. The guide RNAs were ligated into plasmid backbone (pXPR_003, Addgene) and the sgRNA plasmid along with a plasmid containing Cas9 (pLX_311-Cas9, Addgene) were transfected into iPSCs using Lipofectamine 3000. Gene editing was confirmed using the GeneArt Genomic Cleavage Detection Kit and the target locus was amplified with PCR and sent for sequencing. Decreased *INPP5D* expression was confirmed using qPCR and western blotting for mRNA and protein levels following successful differentiation to iMGLs. All guide RNA and primer sequences are detailed in **Supplementary Table 4**.

## ACKNOWLEDGEMENTS

We thank D. Selkoe for his advice on the project; the NeuroTechnology Studio at Brigham and Women’s Hospital for providing LSM710 and Chromium 10x instrument access and consultation on data acquisition and data analysis; A. Stern for postmortem human brain samples for human brain immunostaining; L. Liu for advice on human brain immunostaining; C. Muratore (iPSC NeuroHub at the Ann Romney Center for Neurologic Diseases) for guidance on iPSC-derived neuron cultures; D. Duong (Emory Integrated Proteomics Core at Emory University School) for technical assistance with TMT-MS; Harvard Stem Cell Institute for the CX3CR1-GFP iPSC line; I. Chiu, B. Stevens, and J. El Khoury for their Dissertation Advisory Committee guidance and manuscript feedback, O. Butovsky for the P2RY12 antibody; and to members of the Young-Pearse lab for their critical reading and manuscript feedback. This work was supported by NIH grants F31AG063398, P30AG10161, P30AG72975, R01AG15819, R01AG17917, U01AG46152, U01AG61356, and R01AG063398.

## DATA AND CODE AVAILABILITY

RNAseq and proteomic data sets from iPSC-derivatives and brain tissue are available through the AMP-AD Knowledge Portal (https://doi.org/10.7303/syn25169976). ROSMAP cohort data can be requested at https://www.radc.rush.edu. All original code has been deposited at Zenodo and is publicly available as of the date of publication.

## EXTENDED DATA FIGURE TITLES AND LEGENDS

**Extended Data Fig. 1:**
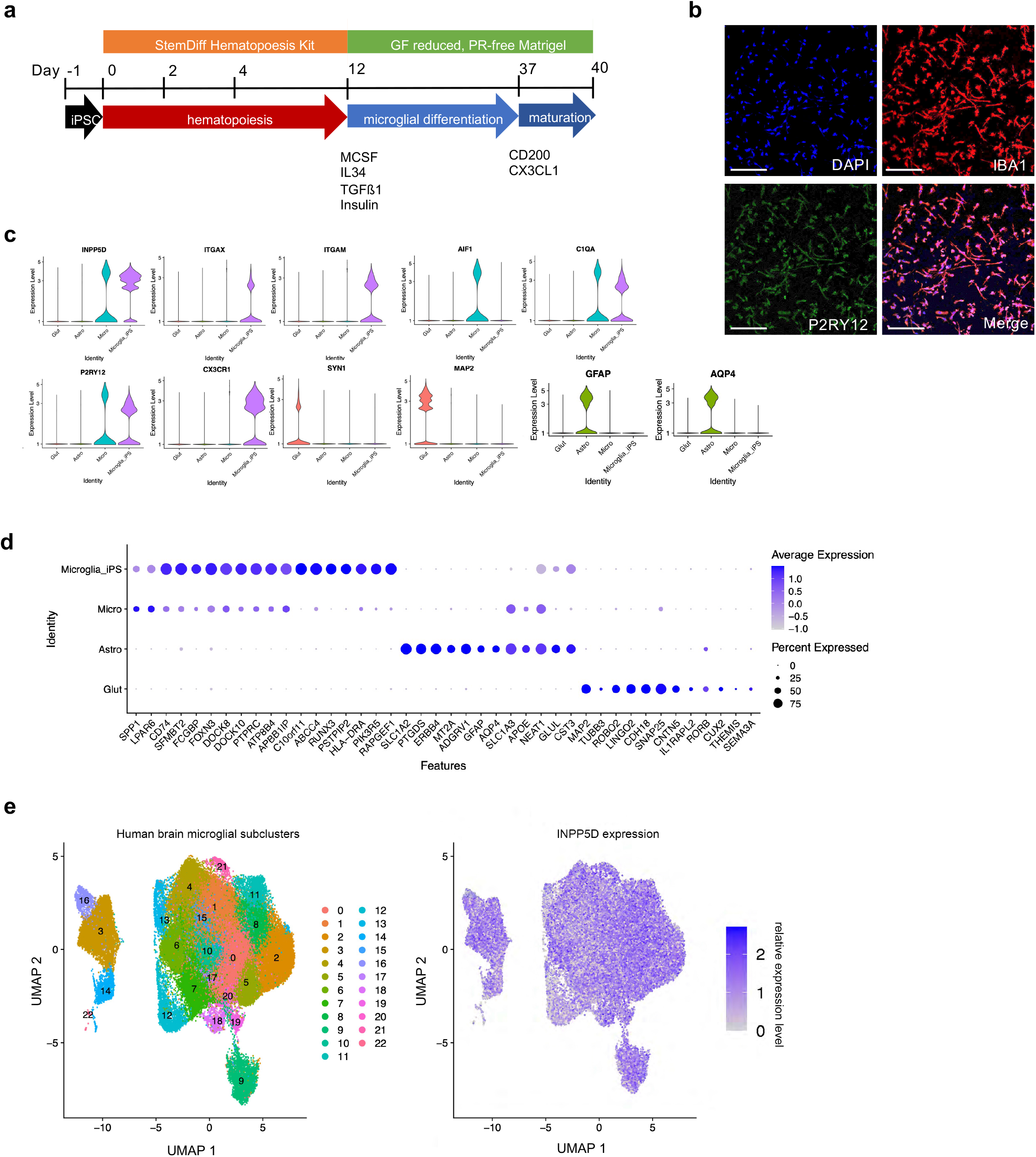
Characterization of iPSC-derived microglia. **a**. iMGL differentiation timeline^26,27^. **b**. Immunostaining of iMGLs for P2RY12 and IBA1. Scale bars = 200 um. **c**. Violin plots of cell fate marker genes for microglia, neurons, and astrocytes from single nucleus sequencing of iMGLs (Microglia_iPS), and glutamatergic neurons (Glut), astrocytes (Astro), and microglia (Micro) from dorsolateral prefrontal cortex (dlpfc) from 12 human postmortem brain samples^23^ and iMGLs. **d**. Average expression of cell fate markers genes across the iMGLs and microglia, astrocytes, and glutamatergic neurons from postmortem brain samples. **e**. Human brain microglia subclusters from the snRNAseq (left) and the relative INPP5D expression across the subclusters (right).

**Extended Data Fig. 2:**
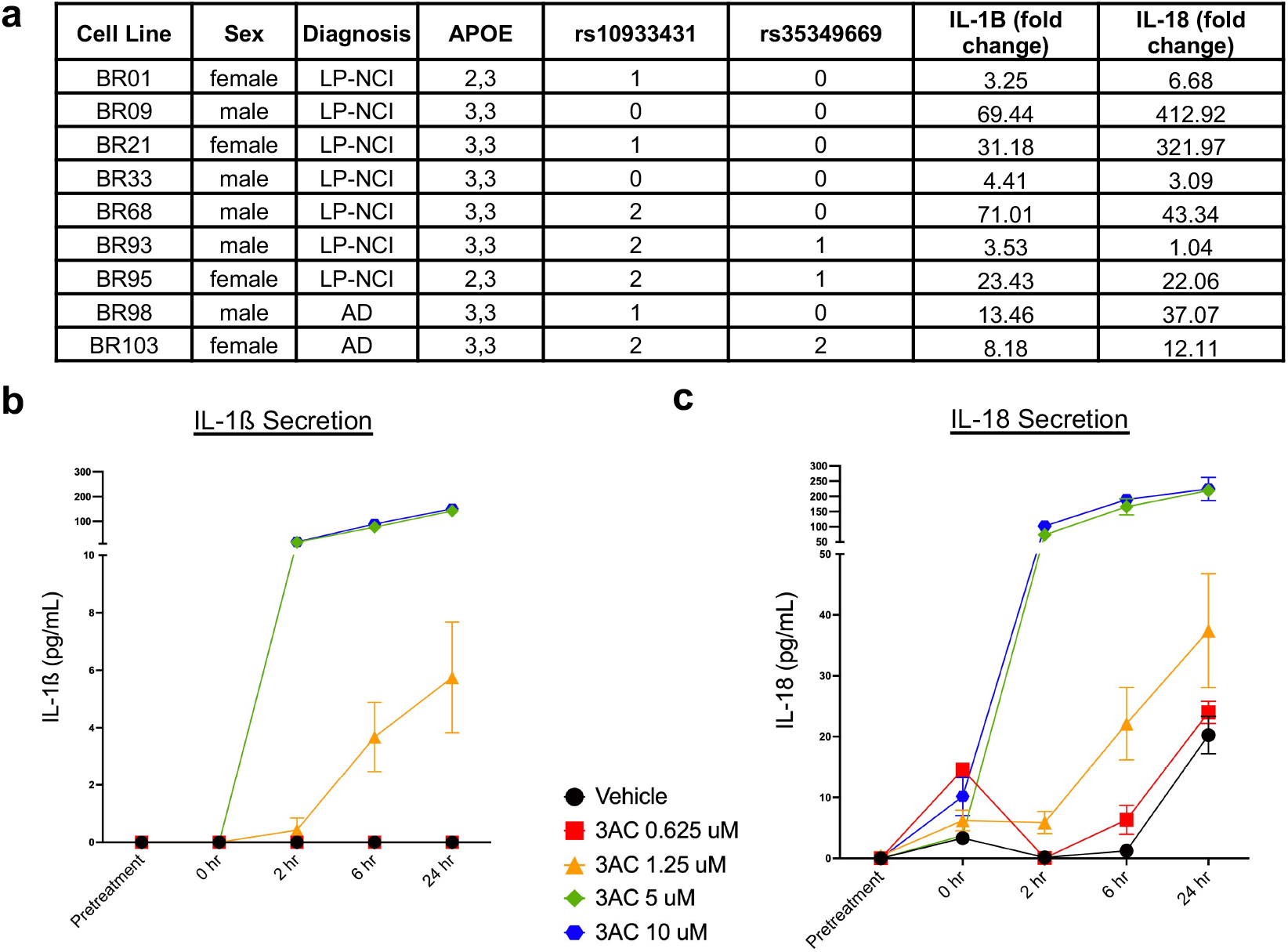
3AC treatment induces IL-1ß and IL-18 secretion in iMGLs across multiple genetic backgrounds. **a**. A table of data regarding the 8 iPSC lines utilized, and the measured fold change of IL-1ß and IL-18 after 3AC (1.25 uM 3AC, 6 hours) treatment. **b-c**. IL-1ß and IL-18 measured at various timepoints (0 hour, 2 hour, 6 hour, 24 hour) of BR33 iMGLs treated with increasing concentration of 3AC (Vehicle, 0.625 uM, 1.25 uM, 5 uM, 10 uM 3AC). n=3-4 wells per condition.

**Extended Data Fig. 3:**
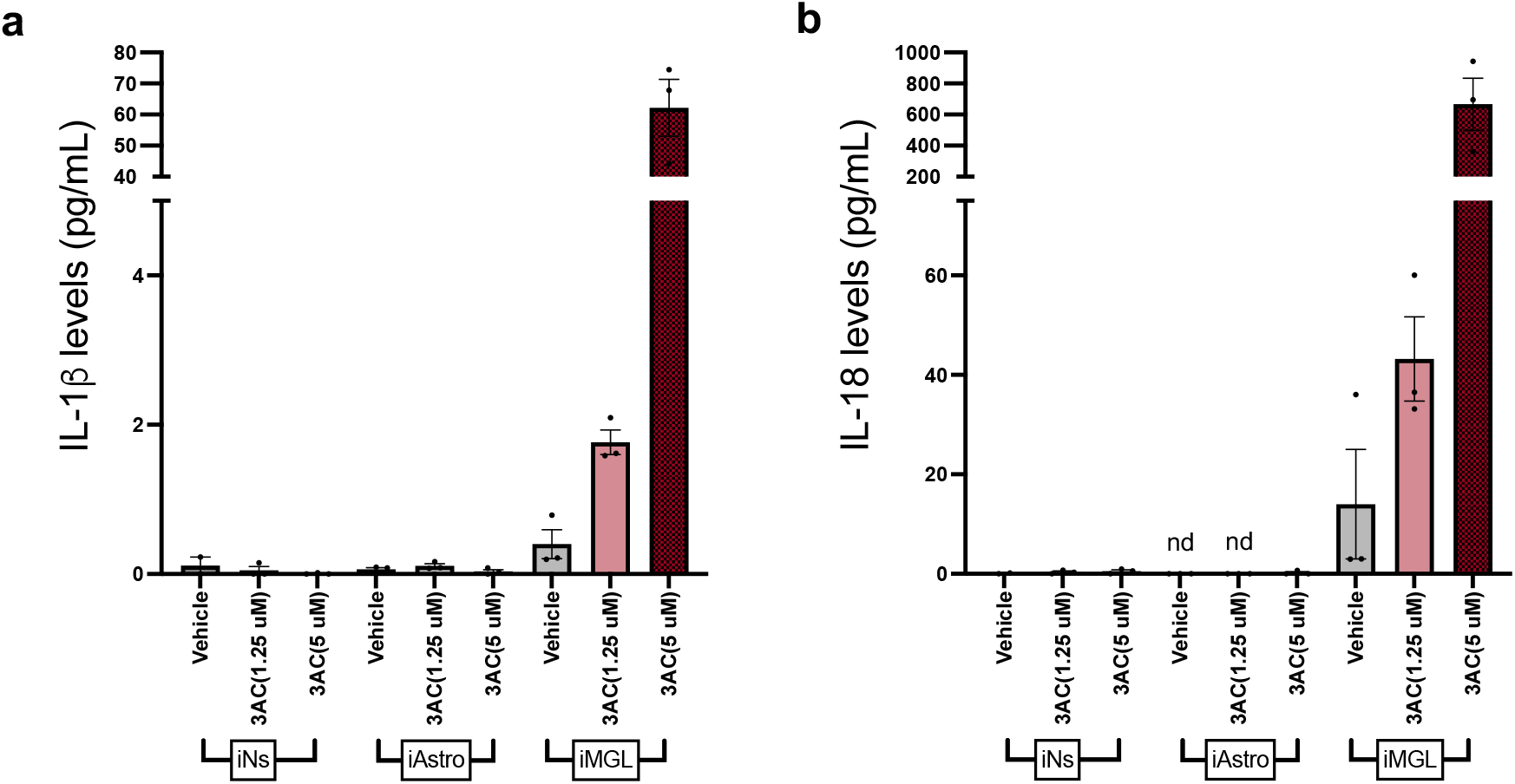
Secreted IL-1ß and IL-18 measured from iPSC-derived brain cell types (iNs, iAs, and iMGLs) treated with 3AC (1.25 uM or 5 uM, 6 hours). IL-1ß and IL-18 were measured using MSD ELISA following treatments. All samples were run on one ELISA plate to compare across cell type conditions. n=3 wells per condition. Mean +/-SEM. nd = not detected, all samples were below detection range.

**Extended Data Fig. 4:**
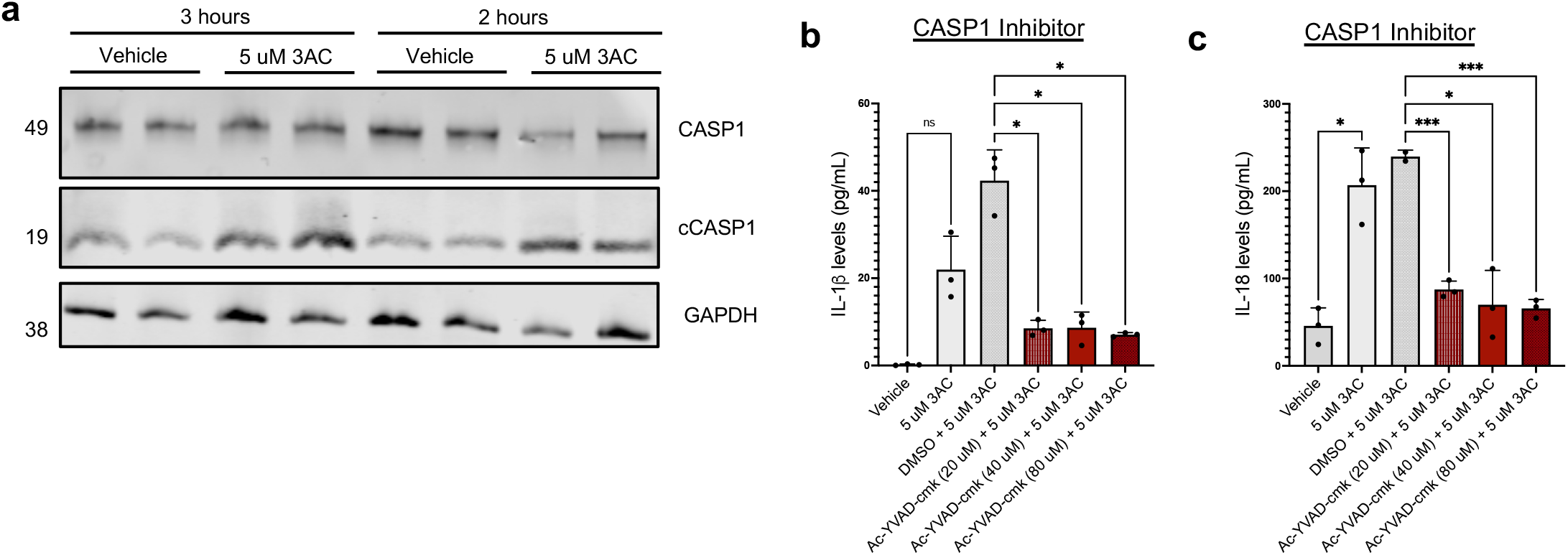
Cleaved caspase 1 increases following INPP5D inhibition. **a**. iMGLs were treated with either vehicle (ethanol) or 3AC (5 uM) for 3 or 2 hours and harvested to collect protein. Caspase 1 and cleaved caspase 1 levels were assayed via western blotting. **b-c**. Treatment of iMGLs with either vehicle (ethanol) or 5 uM 3AC with either DMSO or Ac-YVAD-cmk (20 uM, 40 uM, 80 uM) pre-treatment for 1 hour. Levels of secreted IL-1ß and IL-18 were measured using an MSD ELISA. n = 3 wells per condition. Mean +/-SEM. One-way ANOVA with Sidak’s multiple comparison test. For b-c: ns = not significant, p>0.05, *p < 0.05, *** p < 0.001.

**Extended Data Fig. 5.**
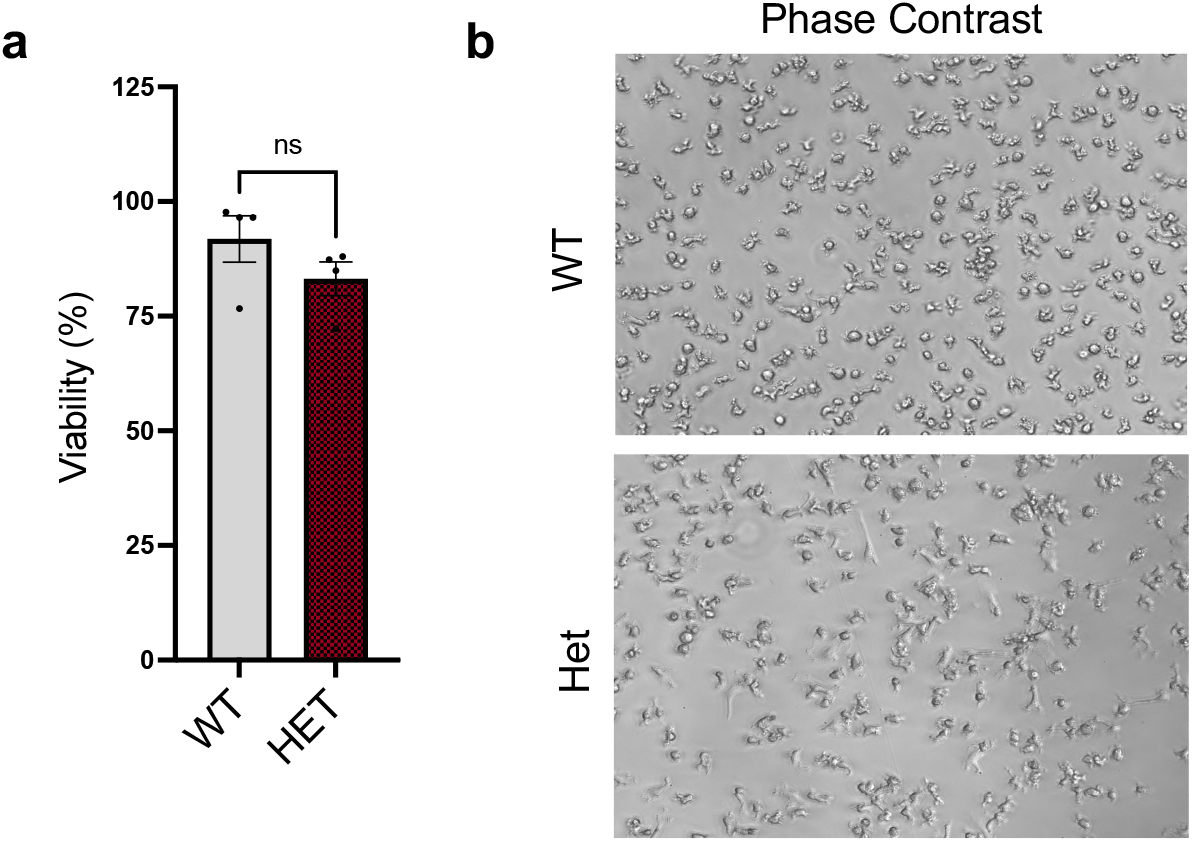
**a**. Viability percentage of the INPP5D WT and HET cells. Viability was determined by measuring the level of LDH secretion normalized to total measure of LDH following cell lysis. n=3. Mean +/-SEM. Mann-Whitney test. ns = no significance (p>0.05). **b**. Representative phase contrast images of INPP5D WT and HET iMGLs.

**Extended Data Fig. 6.**
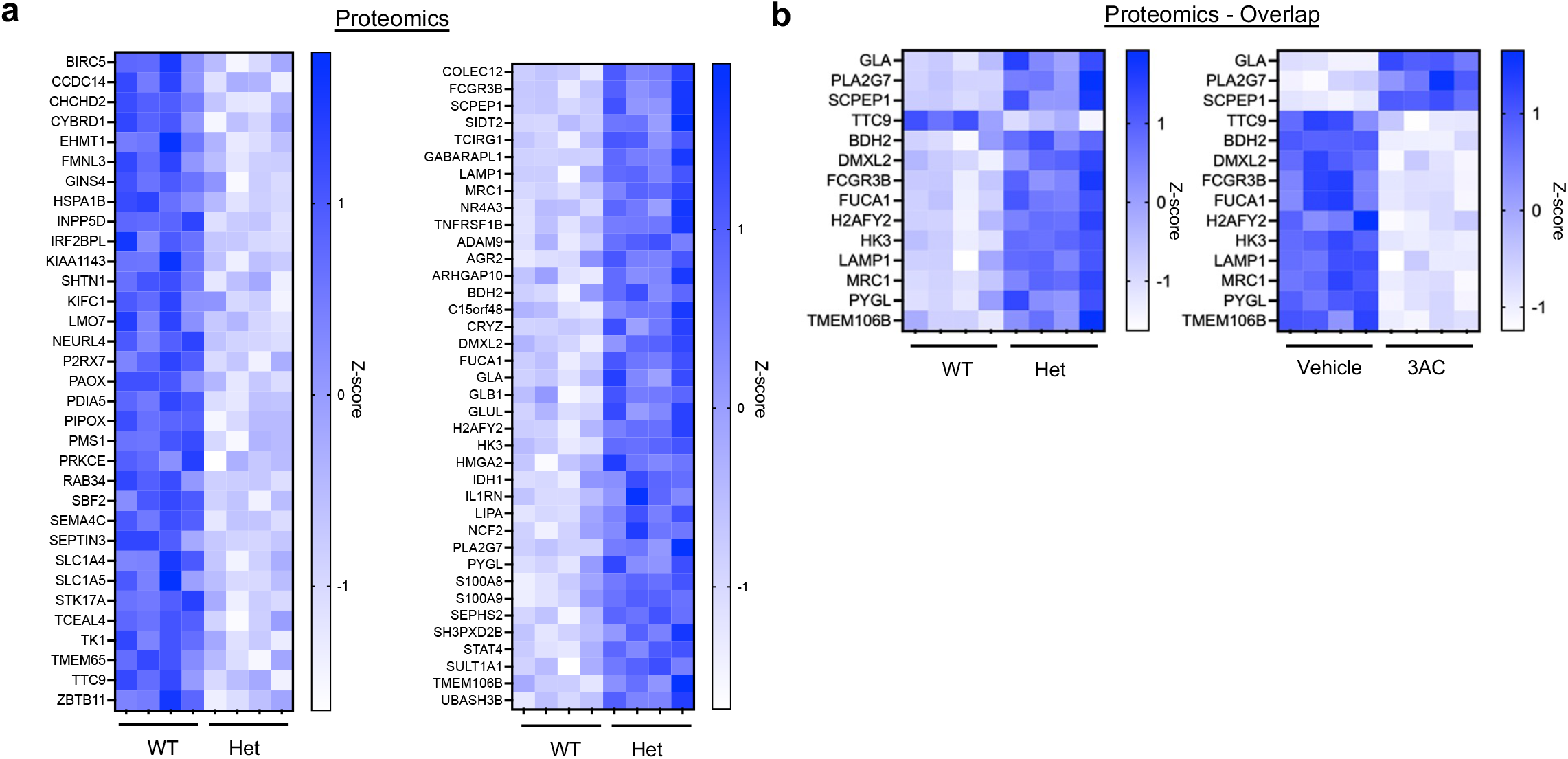
Proteomic changes between INPP5D WT and HET iMGLs. **a**. Heatmap of relative expression of all the differentially expressed proteins comparing INPP5D WT and HET iMGLs (BH FDR<0.05). **b**. Heatmap of relative expression of overlapping differentially expressed proteins between chronic decrease of INPP5D activity (WT and HET) and acute decrease of INPP5D activity (vehicle vs 3AC). Of note, many DEPs are in the opposite direction between acute and chronic INPP5D reduction, suggesting a potential feedback mechanism following 3AC treatment.

